# On-chip large-scale-integration and 2D collective modes of genetically programmed artificial cells

**DOI:** 10.1101/2024.01.10.575012

**Authors:** Joshua Ricouvier, Pavel Mostov, Omer Shabtai, Alexandra Tayar, Eyal Karzbrun, Aset Khakimzhan, Vincent Noireaux, Shirley Shulman Daube, Roy Bar-Ziv

## Abstract

The on-chip large-scale-integration of genetically programmed artificial cells capable of exhibiting collective modes is an important goal for fundamental research and technology. Here, we report assembly of a 2D layout of 1024 monolithic DNA compartments as artificial cells on a 5-millimeter square silicon chip. Homeostatic cell-free protein synthesis reactions driven by genetic circuits occur inside the compartments. We created a reaction-diffusion system with a 30x30 square lattice of artificial cells interconnected by thin capillaries for diffusion of products. Driving the system by a genetic oscillator revealed emergent collective modes of synchrony and propagating phase waves in 2D, with dynamics controlled by geometry. This demonstrates a class of nonequilibrium autonomous systems, where chemical energy consumed to make proteins induces 2D collective patterns of gene expression on multicellular scales, with applications in biological computing, sensing, and materials synthesis.

---

In the 1970s, miniaturization and fabrication of monolithic 2D layers on silicon chips enabled large-scale integration (LSI) of thousands of individual interacting components onto an area less than 5mm square, leading to the development of integrated circuits and the microprocessor. Similarly, in the early 2000s microfluidic LSI technology was made possible by fabricating a layer of soft monolithic microvalves to control thousands of fluid chambers in a multiplexed and addressable manner (*1*). By analogy, on-chip LSI of artificial cells programmed by cell-free protein synthesis holds the potential to create synthetic biological devices capable of autonomous collective behavior and biological computation across scales much larger than a cell. The development of cell-free genetic circuits (*2–5*) has pushed forward the assembly of minimal cell-models in embodiments such as liposomes, emulsions, coacervates, and solid-state compartments (*6–13*), with work toward cell-cell communication showing examples of front propagation, synchronization, light activation, and signaling (*14–20*).

We developed the solid-state compartments as a step toward “artificial cells” on a chip (*9*). A series of miniaturized structures carved on a chip and DNA-driven reactions, with materials diffusing into and between the compartments, recreated steady-state and oscillating protein expression patterns. So far, however, the design has been topologically limited to 1D chains of communicating artificial cells (*15, 19*). As in many areas of science and technology, the transition from 1D to 2D artificial cell systems would expand the possibilities, and reveal phenomenology of collective modes owing to the increase in connectivity and possible arrangements. Here, we transitioned from 1D to 2D on-chip artificial cells by carving feeding wells that traverse the chip from side to side, creating a perforated membrane-like structure. Using the third dimension to connect every cell with the outside environment for exchange of nutrients solves the topology problem, enabling 2D layouts of interconnected artificial cells.

We first present the LSI of 1024 disconnected artificial cells on an area of 5x5mm2, and then show an interconnected square lattice implementing a reaction-diffusion system. We programmed the connected lattice by a genetic circuit of a nonlinear oscillator (*9, 19*), and discovered emergent collective modes of synchrony and wave propagation. The dynamics of the system, including effective protein lifetime and diffusion constant, strength of intercellular coupling, and propagation speed are all controlled by geometry. This opens a class of nonequilibrium programmable active matter cellular systems (*21*), where chemical energy dissipated in protein synthesis leads to spatiotemporal patterns of information encoded in biomolecular concentrations.

## A 3D architecture for LSI of on-chip monolithic artificial cells in 2D

We fabricated a square lattice of 32x32 independent artificial cells (unit length a = 150μm) on a silicon wafer with protein synthesis compartments at the front side, and feeding channels at the back (Fig 1). The compartments are shaped as thin pancakes (radius R=35μm; thickness h=3μm), with narrow capillaries (width w=8μm; length L=80μm) connected to perpendicular wells traversing the chip from side to side (radius R_W_=14μm; length H=250μm). The back side includes 32 parallel microfluidic channels, each aligned with a row of wells (fig. S1). Linear double-stranded DNA (dsDNA) brush (20x20μm2) are immobilized on the surface of every compartment, each addressable and programmable by a different set of gene constructs(fig. S2). Protein synthesis occurs when an E.coli cell-free transcription-translation (TXTL) system (*22, 23*) flowing in the feeding channel diffuses through the well and capillary to initiate protein synthesis at the DNA brush source. In turn, newly made proteins diffuse out of the compartment along the capillary and the well, where they are transported out by the flow at the back side. Each DNA brush co-localizes TXTL machinery and products, creating favorable conditions for gene regulation and multi-component interactions (*27, 28*). With up to 1000 dsDNA/μm^2^, the brush enables control of gene composition, and copy number down to the single gene level where expression is stochastic (*29*)).

**Fig. 1.**
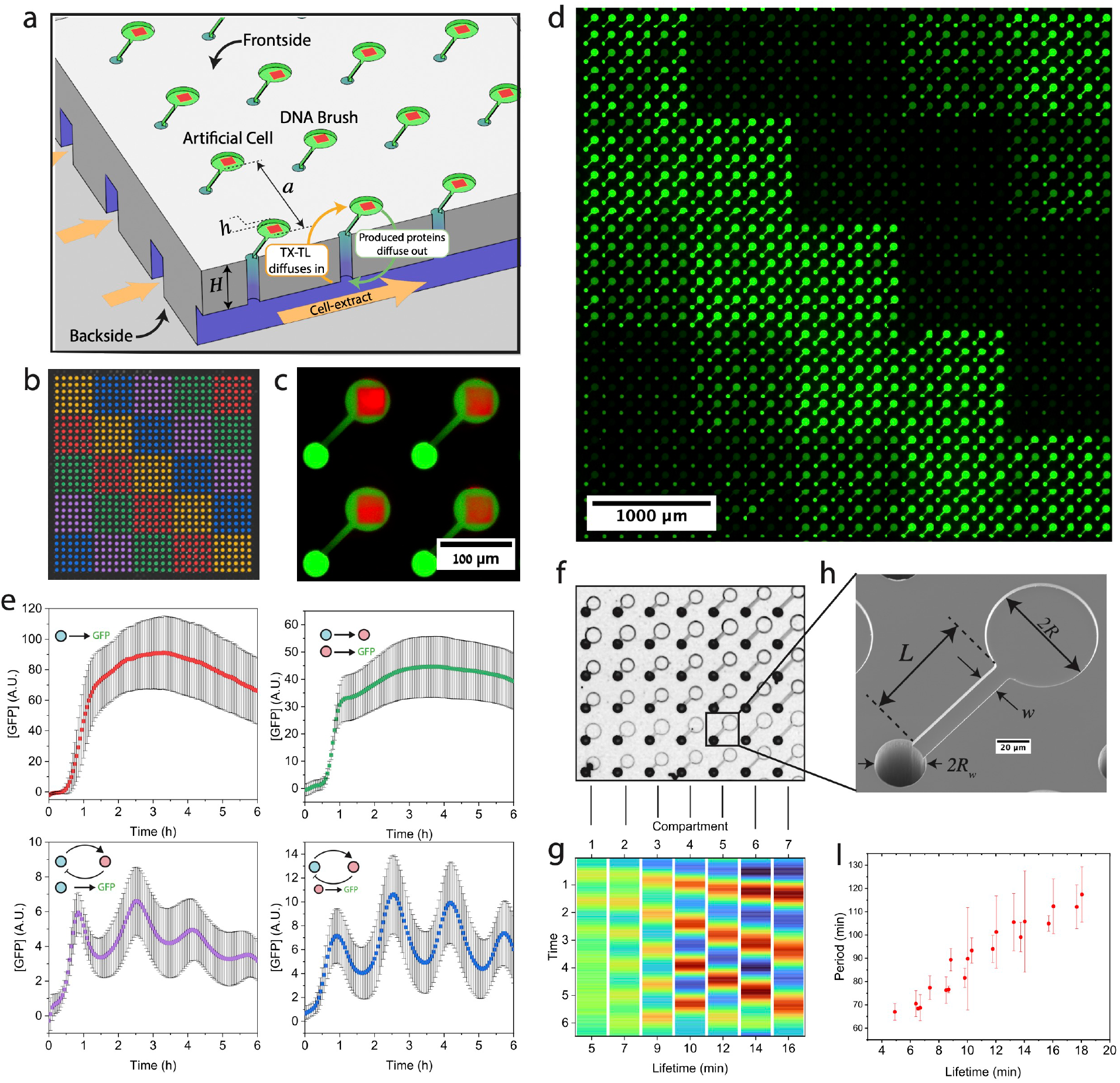
Large scale integration of artificial cells on a chip. A – Scheme of chip. Frontside: A 2D array of artificial cells, each comprising of a disk-shaped compartment carved h = 3μm deep, with immobilized DNA brush, and a thin capillary connecting the compartment to the backside through a well. Backside: microfluidic channels transport E.coli TXTL system diffusing through the well into the compartment to initiate protein synthesis. In turn, newly made proteins diffusing out of the well are flown away so as to create a source-sink and hence steady-state gene expression. B –Allocation of compartments according to gene constructs spotted. Red: P70-*gfp*; green: P70-*σ28*, P28-*gfp*; magenta: Oscillator reported by P70-*GFP*; blue: Oscillator reported by P28-*gfp*; orange: PT7-*gfp*. Oscillator is a set of five genes: P70-*σ28*, pTET-*Aσ28*, P28-*ci-ssra*, PT7-*clpP*, PT7-*clpX*. C – Fluorescent overlayed images of DNA strands (red, Alexa 647) and cell-free expressed GFP (green). D – Fluorescent image of GFP expression according to map in B at t=3.35h. The chip contained 900 identical compartments placed in a square lattice. E – Expression profiles according to B. F– Bright field image of a chip of compartments with different lifetimes (5-19 min). G – Expression of genetic oscillator in corresponding compartments. H – Scanning electron microscope image of a single compartment, scale bar 10μm. I – The period of the oscillator as a function of the compartment lifetime.Fig. 2. You can place graphics in-line above each caption. Please do not use text boxes to arrange figures. High-resolution (preferably editable PDF or Adobe Illustrator format) figure files will be requested following review.

The scenario of a localized protein source at the DNA brush and a sink at the bottom of the well, implies that the protein concentration will reach steady-state when the rate of synthesis balances the rate of dilution out by diffusion. The solution of the diffusion equation in the compartment is a piece-wise linear concentration gradient, dropping from the source along the capillary, and then along the well to the exit point sink, where the concentration vanishes due to the flow (SI; fig. S3). Theoretically, the system reaches steady-state within an effective lifetime that is set by geometry, *τ* ≈ *τ*_0_(1 + Δ*L*/*L* ), where *τ*_0_ = *πR*^2^*L*/*D*_0_*w* = 16 min is the compartment lifetime without the well, and *D*_0_ ≈ 40μ*m*^2^/*s* is the typical diffusion constant of newly made proteins (*9*). The contribution of the well is to effectively elongate the capillary by a relative factor 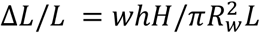, which is only about 12*c* because the capillary has comparable length to the well, *H* ≈ *L*, but smaller cross section, 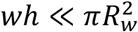 (fig. S3). This implies that protein concentration in steady-state should scale with *τ*_0_ (fig S4).

To prove experimentally that the wide well has negligible contribution to the lifetime of the artificial cells, we set out to measure protein synthesis dynamics and test the source-sink model and scaling of steady-state with lifetime. The 2D approach provides simple means to test several different DNA constructs and genetic programs in parallel on a single array, with GFP as a reporter in all of them, for accurate comparison: an unregulated strong promoter, a transcriptional cascade, and a nonlinear oscillator constructed by an activator-repressor feedback circuit (Fig. 1d-e, fig. S4). The data showed an increase in protein synthesis for about 1hr, followed by a transition to a steady concentration for the unregulated and cascade, and to oscillations for the activator-repressor feedback circuit, which is consistent with a source-sink scenario. As expected, the spatial concentration profile dropped linearly along the capillary from the DNA brush to the center of the well, extrapolating to zero at a point located only Δ*L*/*L* ≈ 18*%* down the well, implying that the sink is not altered much by the presence of the well (Fig S3). We then tested the influence of geometry on the synthesis dynamics using a chip of varying capillary length and compartment radius, and observed that steady-state concentrations, oscillator amplitude, and period scale with *τ*_0_, thereby validating the source-sink model (Fig. 1f,g,i, fig. S4).

## Programmable reaction-diffusion on a square lattice of artificial cells

To transition from isolated artificial cells to a coupled system, we fabricated a device having a square lattice of cells interconnected by thin capillaries of width w_c_ = 8 μm, and length L_c_ = a − 2R = p5 μm. Proteins synthesized at a given source diffuse out to the well and into its four neighboring cells, where they recurrently dilute out through the respective additional wells. Therefore, by the same source-sink model, the concentration profile diffusing out of the source is expected to drop linearly at every step on the lattice. The geometrical parameter that characterizes the coupling strength between neighboring cells is *β* = *Lw*_*c*_/*L*_*c*_*w*.

To validate this picture, we measured the dynamics of protein synthesis and diffusion in a lattice of *β* = 0.9 by immobilizing a few isolated sources of proteins expressed under a strong promoter, surrounded by empty compartments, with a few uncoupled sources, and single sources of oscillators, on the same chip for reference (Fig. 2). The dynamics of the isolated sources was not stable in time due to the dilution out to the neighboring cells, in contrast to the dynamics within a 30x30 coupled lattice of sources, which remained stable in time, as with uncoupled compartments (fig S5). Consistent with diffusion from the source, the spatial profile of the source dropped linearly along the capillaries at every step, and decayed to background levels three cells away. The profile was essentially symmetric in all directions, except for a minor drift of 6% along the direction of the underlying flow channel at the bottom layer (SI). From the space-time plot of source expression dynamics, we extracted an effective diffusion constant of D_eff_ = 2p ± 2 μm^2^/s. We also measured the expression profile of single oscillator sources and the diffusion of the reporter to its neighbors, estimating a similar diffusion constant, and a short delay between the oscillating peaks in neighboring cells of Δt = 9 min (fig. S5).

**Fig. 2.**
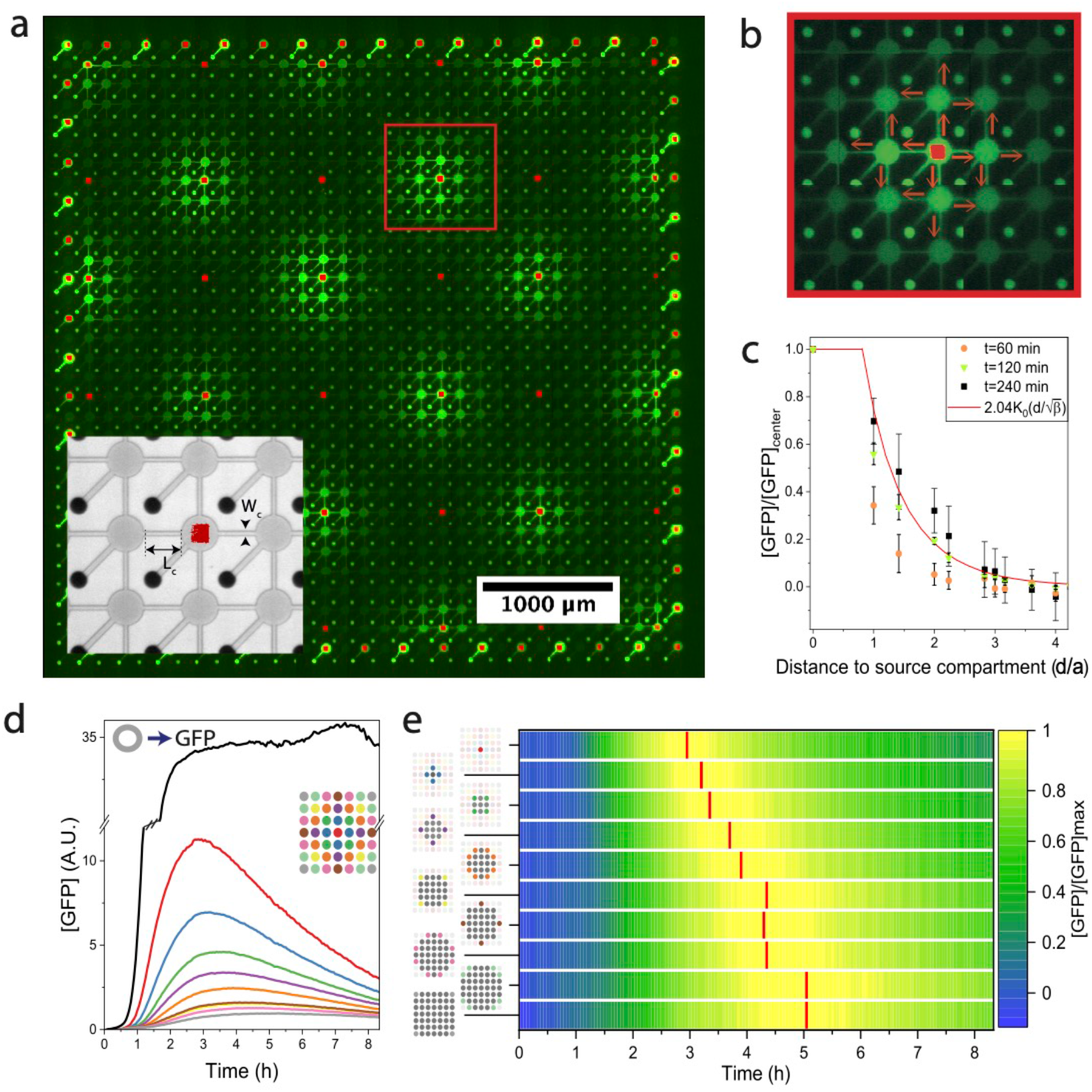
Diffusion from a single source in a coupled array of compartments. A – A chip of 30x30 connected compartments surrounded by isolated ones along the boundaries. Inset: bright field image and fluorescent image of DNA. Connecting capillaries length *L*_*c*_ = 70*μm*, and width *w*_*c*_ = 12*μm*, and coupling strength 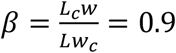. Fluorescent image of GFP expression from a dilute array of single sources (P70-GFP; square DNA brushes labeled in red; P70-gfp, 16 bright regions; oscillator circuit, 18 dim regions (reported in Fig S5)). Scale bar 1000μm. B – Close-up of one source expressing GFP and diffusing into neighboring compartments. C – The spatial profile at t=60min (orange disk), t=120 min (green triangle) and t= 240min (black square) as a function of the Euclidean distance d of neighbors. The solid red line is the analytical solution 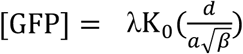. D – The temporal expression profile of the source (red) and profiles of diffused GFP into neighbors (distance designated by color) with a reference of an uncoupled compartment (black). The 1st neighboring compartments are averaged and reported by the blue line. Green, purple, orange, yellow, brown, pink, light green and gray, report on the nth neighboring compartments, respectively. E – Plot of expression profile as a function of time and cumulated compartment area from source, with maximal GFP concentrations designated in red. A linear fit to the maxima yields an effective diffusion constant of D_eff_ = 27 ± 2 μm^2^/s

The transition from a single source to coupled system of sources on the lattice implies a discretized 2D reaction-diffusion equation for the concentration of proteins *P*_*i,j*_ at coordinates (i,j) (SI):

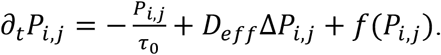

Here, Δ*P*_*i,j*_ is the discrete Laplacian of the protein concentration, 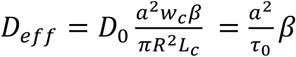 is the effective coefficient diffusion on the lattice. The diffusion within the array is solely defined by the geometry, and the coupling strength beta plays a predominant role *β* = *τ*_0_/*τ*_*c*_, with *τ*_*c*_ = *πR*^2^*L*_*c*_/*D*_0_*w*_*c*_ the lifetime for diluting out into neighboring cells. The steady-state solution of a single source in a continuum approximation, is 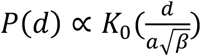, *K*_0_ the modified Bessel function (*d* is the Euclidian distance from the source), which fits the data (Fig. 2). The function *f*(*P*_*i,j*_) is programmed by the genetic code embedded in each compartment. With feedback circuits such as the genetic oscillator, this leads to nonlinear collective dynamics, which generally support propagating waves (*24*) with a lower bound on the speed set by an intrinsic value that is controlled by the geometry of the chip:

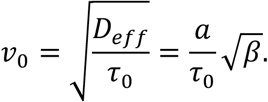

## Synchrony and phase waves in a lattice of coupled genetic oscillators

To test whether the LSI of coupled artificial cells leads to emergent collective behavior, we immobilized the genetic oscillator circuit on 30x30 cells of a connected lattice (*β* = 1.1), and observed the collective dynamics (Fig. 3). The oscillations of 900 cells began in unison with a mean period of T = 119 ± 3 min. In time, we observed large-scale spatial variations, dephasing from the perfectly synchronized collective oscillations (*25, 26*), and waves propagating inward from the boundaries, (Fig. 3d), as shown in a window of 10x10 cells (Fig. 3e). A space-time plot of a line of 10 consecutive cells inside the lattice shows continuity of phases as a signature of synchrony, in contrast to a line of uncoupled oscillators (*β* = 0, fig. S7). Past the third oscillation, the vertical lines of peaks deflected to a finite slope characteristic of a phase wave with a measurable speed of 0.6 ± 0.1μm/s, which is of the same order and a bit faster than 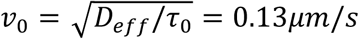. To further characterize the collective modes we computed the power spectrum of lattice dynamics as a function of wave vector and angular frequency (k, ω) (Fig. 3h). For the main mode at the natural frequency of the oscillator, ω = 2π/T, the amplitude drops with wave vector, and vanishes for k = 2π/λ corresponding to a wavelength of *λ* ≈ 8 cells. These modes correspond to phase velocities v_p_ = ω/k in the range 0.19 < v_p_ < 0.6μm/s.

**Fig. 3:**
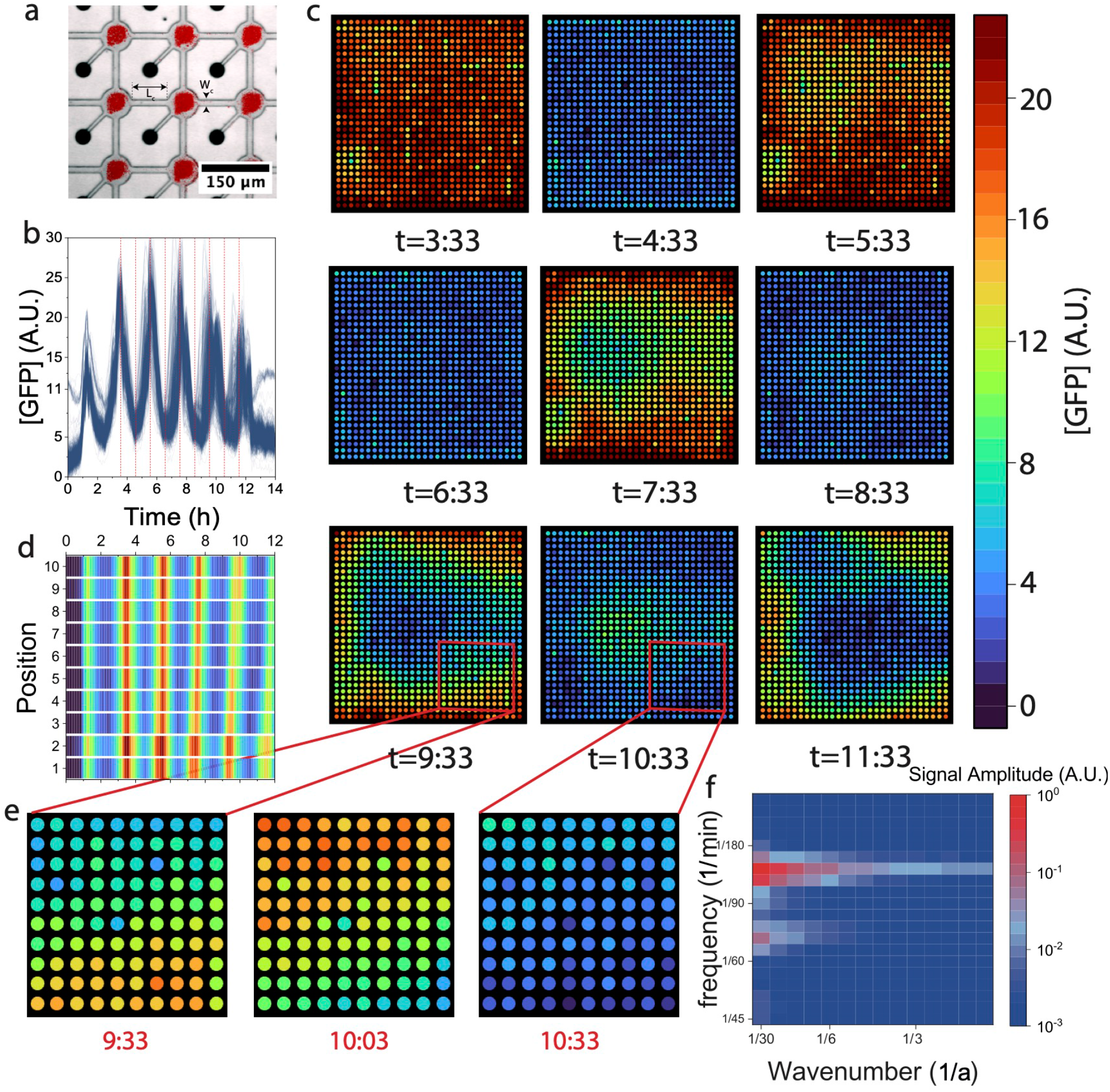
Collective patterns of a coupled array of genetic oscillators. A – Overlay of brightfield and DNA brush fluorescence (red squares) in a coupled array of 30x30 compartments with couple strength *β* = 1.1. B – GFP expression dynamics of all 900 oscillating compartments, with average period of T=120 min. C – Spatial expression profiles of entire 30x30 array of compartments at nine time points, in hr:min as denoted. D Space-time pattern for a line of ten compartment orthogonal to the wave. E. Close-up of the emerging propagating wave at three time points. F– Power spectrum of the collective oscillations as a function of frequency and wavenumber.

To further characterize the collective behavior in the coupled oscillating lattice of cells, we extracted the instantaneous phase of every cell, ϕ(t) (Fig. 4a). The standard deviation of all 900 oscillator phases increased in time (Fig. 4a inset), consistent with the loss of perfect synchrony due to the emergence of phase waves. We plotted the 2D isophase function of the lattice just after the fifth peak (Fig. 4b), which revealed a pattern showing that oscillators at the boundaries were in advance, whereas those in the center were retarded relative to the mean. We next calculated the gradient vector field of the isophase function, which reports on the phase waves (Fig. 4c). Regions with zero gradient indicate synchrony, whereas phase waves move along finite gradients at velocity inversely proportional to the gradient field magnitude. The gradient field revealed concentric phase waves propagating from the boundaries inward into the center of the lattice, implying spatial cooperativity of all the artificial cells.

**Fig. 4:**
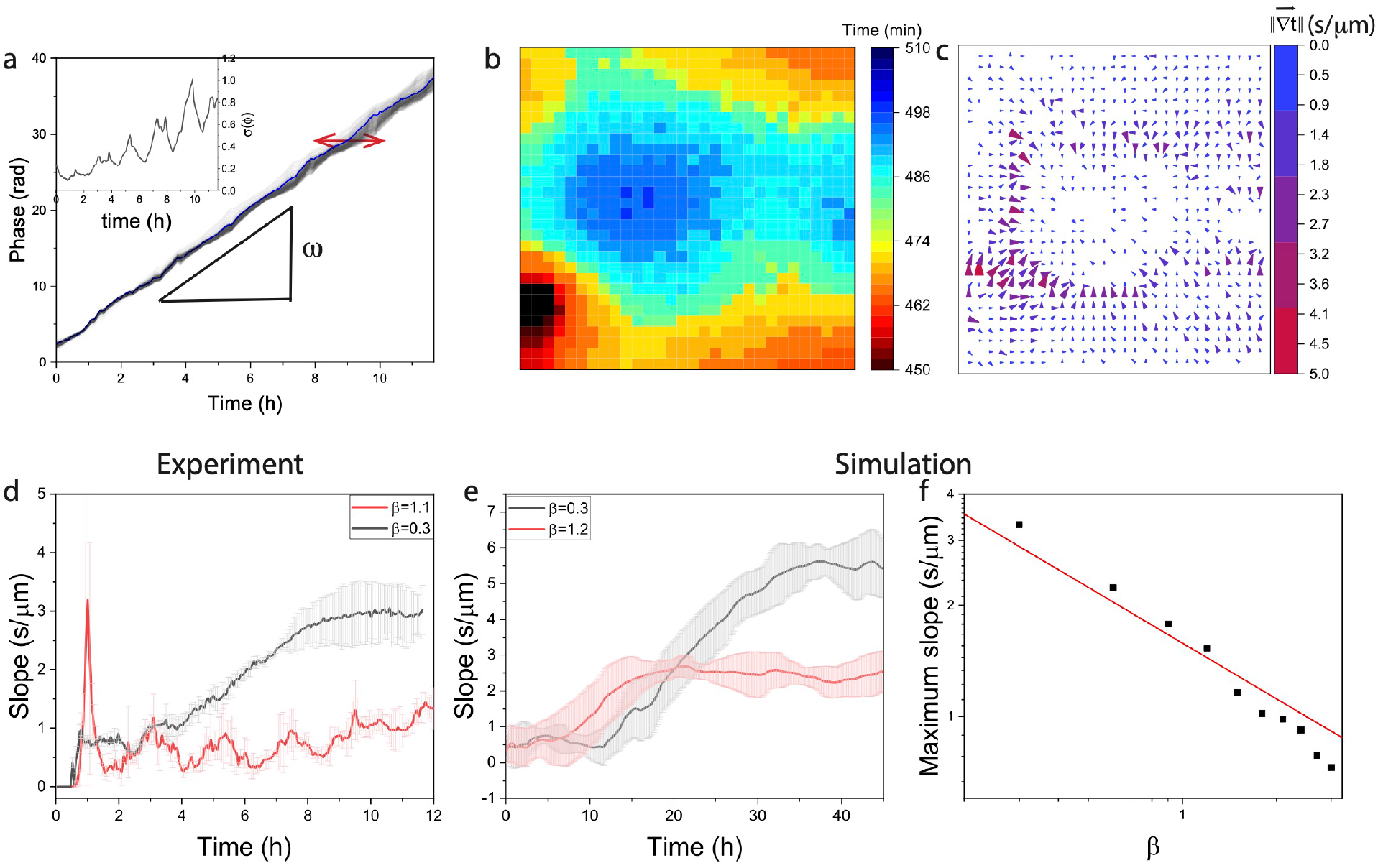
Phase waves and the emergence of coordination across the array. A – The continuous phase of the oscillators as a function of time for all 30x30 coupled compartments (Figure 3), with slope of the mean phase (solid blue line) as the angular frequency of the oscillators. Inset – Standard deviation of the phase. B - Isophase surface for *ϕ* = 28 rad (after the 5th oscillation). C – 2D gradient vector field of the isophase surface. Phase waves propagate along the direction of the vector field, with a speed inversely proportional to the gradient magnitude. Defects, corners and edges generated phase waves converging to the center of the chip. D – Gradient (slope) (projected on the x-axis) along time for the array with a coupling strength of *β* = 1.1 and *β* = 0.3. The slope is inversely proportional to the wave speed E – Simulation of the corresponding arrays of coupled oscillators with an emerging wave propagating within the system. F – Simulation of the maximum slope plotted for different values of *β* (scatter), referenced against a power law with exponent of −1/2 (red).

Finally, to test how the phase wave speed depends on geometry, specifically whether it scales with coupling strength, 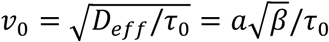, we assembled a lattice of connected cells having similar lifetime, but weaker coupling strength, *β* = 0.3. This lattice was indeed “softer” with a profile of isolated protein sources that decayed on a scale of just one cell (fig. S6, S7), and a correlation length of *ξ* = 1.6*a*, compared to the more “rigid” *β* = 1.1 lattice for which *ξ* = 3.9*a* (fig S8). We then programmed the entire 30x30 lattice by the genetic oscillator circuit, and measured the mean slope in time, which increased after 10 hours to a maximal value corresponding to a minimal phase wave speed value of *v* = 0.33 μm/s for *β* = 0.3, compared to *v* = 0.66 μm/s for *β* = 1.1. We confirmed this result by simulating the reaction-diffusion equations of the oscillator (SI, Fig 4e,f), and for several values of *β*, and observed that the lower limit of phase waves speed scales with 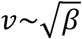. This is in agreement with a broad class of nonlinear reaction-diffusion systems that support propagating fronts, and demonstrates that the physical dimensions of the artificial cells and their coupling determine the collective dynamics of the lattice.

## Discussion

LSI of genetically programmed artificial cells on a chip is inspired by the miniaturization in electronics and by multicellularity in biology. Packing independent artificial cells in 2D, each with a lifetime set by the dimensions of the compartment, and programmed by genetic circuits, enables measuring thousands of gene-expression reactions in steady-state. This will inform on biological design principles with applications in synthetic biology and biotechnology. The transition from a single compartment to a coupled array exhibiting collective dynamics leads to spatiotemporal patterns of information encoded by biomolecular concentrations as in morphogenesis (*30, 31*). The synchronization and phase waves of genetic oscillators are an example of the emergent complexity that is achievable by combining genetic programming with a geometrically controlled reaction-diffusion scenario (*32, 33*), each cell programmable by its own DNA blueprint. Finally, our approach is readily scalable to much larger arrays in 2D and is amenable to further miniaturization by standard microfabrication.

## Acknowledgments

We acknowledge funding from the Israel Science Foundation (R.B.Z. and S.S.D., grant no. 2723/19), the United States – Israel Binational Science Foundation (R.B.Z. and V.N. grant no. 2018208), the Isak Ferdinand and Dwosia Artmann Research Fund for Biological Physics (R.B.Z), from the Human Frontier Science Program (V.N., grant no. RGP0037/2015) and the Minerva Foundation (R.B.Z. and S.S.D., grant no. 712274). J.R. is grateful to the Azrieli Foundation for the award of an Azrieli Fellowship.

## Author contributions

Conceptualization: JR, AT, EK, SSD, RBZ

Production of cell extract: AK,VN

Experimentations: JR, PM, OS

Writing: JR, SSD, RBZ

## Data and materials availability

Data and code will be deposited on a public repository prior to publication.

## Supplementary Materials

Materials and Methods

Supplementary Text

Figs. S1 to S8

Movies S1 to S2

