## Supplementary material for "On-chip large-scale-integration and 2D collective modes of genetically programmed artificial cells": SI

**The PDF file includes:**

Materials and Methods
Supplementary Text
Figs. S1 to S8
References

**Other Supplementary Materials for this manuscript include the following:**

Movies S1 to S2

### Materials and Methods

Silicon chip fabrication

The silicon chip is fabricated in a three-step process.

The silicon wafer (Double side polished, 4", 0.3 mm thickness, test grade, p-type, University wafer) is patterned using standard photolithography techniques (Photoresists S18XX series and AZ 4562, Microchem) and is etched by an ICP-RIE (Surface Technology Systems). The front side is 3 µm deep and is composed of an array of 30x30 connected or disconnected compartments (20-30 µm radius) array, and communication micro-channel (6-12 µm wide). The second step consists of etching the backside of the wafer to a depth of 100 µm. The photoresist is exposed after backside alignment, developed and etched by a Bosch process (200 cycles). A third step is added to carve the well. After frontside alignment, the wafer is aligned and then patterned, as well as trenches delimiting the chip, are then etched all the way through the wafer by a Bosch process (700 loops). The device is then coated on the frontside with a $SiO_2$ layer of 90 nm by using PECVD.

Chip coating

Chips coated with $SiO_2$ are then incubated with a biocompatible photoactivable polymer which self-assembles at the surface [1]. The biopolymer is composed of a PEG backbone, a trialkosyloxane function at one end (reacts with $SiO_2$ deposited at the surface of the chip) and a Nvoc protected amine at the other end. Chips are incubated at 0.2 mg/mL in toluene for 40 min and then rinsed in toluene.

Exposition of the chip

The chip is then aligned and exposed with a 400 nm LASER (Microwriter - Durham Magneto Optics Ltd). The monolayer of photoactivable reactive amine group receives a dose of 1000mJ/cm$^2$ spatially patterned

Biotinilation of the chip

The chip is then biotinylated by dipping in a biotin-NHS solution in Borate buffer for 15 min. The biotin-NHS is attached only to the photo-activated amine group leaving the surface patterned with biotin.

DNA preparation

DNA-streptavidin (SA) linear conjugated fragments are prepared by Polymerase Chain Reaction (PCR) in KAPA Readymix (Roche). Biotin and Atto 647N are attached to the DNA fragment by using correspondent primers (IDT). PCR products are then cleaned (Promega Wizard PCR cleaning kit) and then conjugated to SA in PBS. The final DNA concentration is 150-300 nM.

DNA deposition

Dense brushes of dsDNA are formed by the deposition of nanoliter droplets on the artificial cell compartment. Droplets of DNA-SA in PBS are spotted individually (GIX Microplotter, Sonoplot and Sciflexarrayer S3 Scienion). Then they are incubated for 2 hours. The chips are finally rinsed in PBS and then water.

Sealing of the device

The chip is sealed at the backside by a PDMS (Sylgard) slab. The inlet and outlet are pierced through the slab and are aligned with the silicon chip. On the frontside, a glass coverslip covered with a 100 µm thick layer of hard PDMS (Gelest) seals the device. Both the PDMS slab and the coverslip are gently pressed on the silicon chip to ensure sealing.

Imaging

The device is then placed in a temperature chamber at 30°C and is connected to a syringe pump (Harvard apparatus) and a cell-free extract container. The latter is kept at 2°C by a Peltier cell throughout the duration of the experiment. The device is then imaged by an inverted microscope (Axiovert Zeiss) equipped with an automated stage and a sensitive wide view camera (Andor).

Cell-free transcription-translation

Cell-free gene expression was carried out using an *E. coli* TXTL system described previously Briefly, *E. coli* cells were grown in a 2xYT medium supplemented with phosphates[2]. Cells were pelleted, washed, and lysed with a cell press. After centrifugation, the supernatant was recovered and preincubated at 37 °C for 80 min. After a second centrifugation step, the supernatant was dialyzed for 3 h at 4 °C. After a final spin-down, the supernatant was aliquoted and stored at −80 °C. The TXTL reactions comprised the cell lysate, the energy and amino acid mixtures, maltodextrin (30 mM) and ribose (30 mM), magnesium (2-5 mM) and potassium (50-100 mM), PEG8000 (3-4%), water and the DNA to be expressed.

### Supplementary Text

#### I. Backside: parallel flow channels for cell-free extract

The backside is carved in the silicon chip and sealed with a slab of PDMS (Sylgard). It is composed of one inlet, microchannels (depth 120µm) and one outlet. Right after the inlet, a filter divides the incoming flow into 32 parallel flows. The 32 channels are parallel and are connected in order to avoid clogging and bubbles. Typically, the flow rate is 20 µL/h. In one single channel, the flow rate is 0.6 µL/h=0.01 µL/min. The typical dimensions of the parallel flow channels are $h_{backside} = 0.120\ mm$, $w_{backside} = 0.050\ mm$ and $L_{backside} = 5\ mm$. Therefore, the typical volume is 0.03 µL and the flow renews the CFE every 3 minutes.

II. <u>1D Diffusion between the flow-channel and the compartment, Equivalent channel</u>

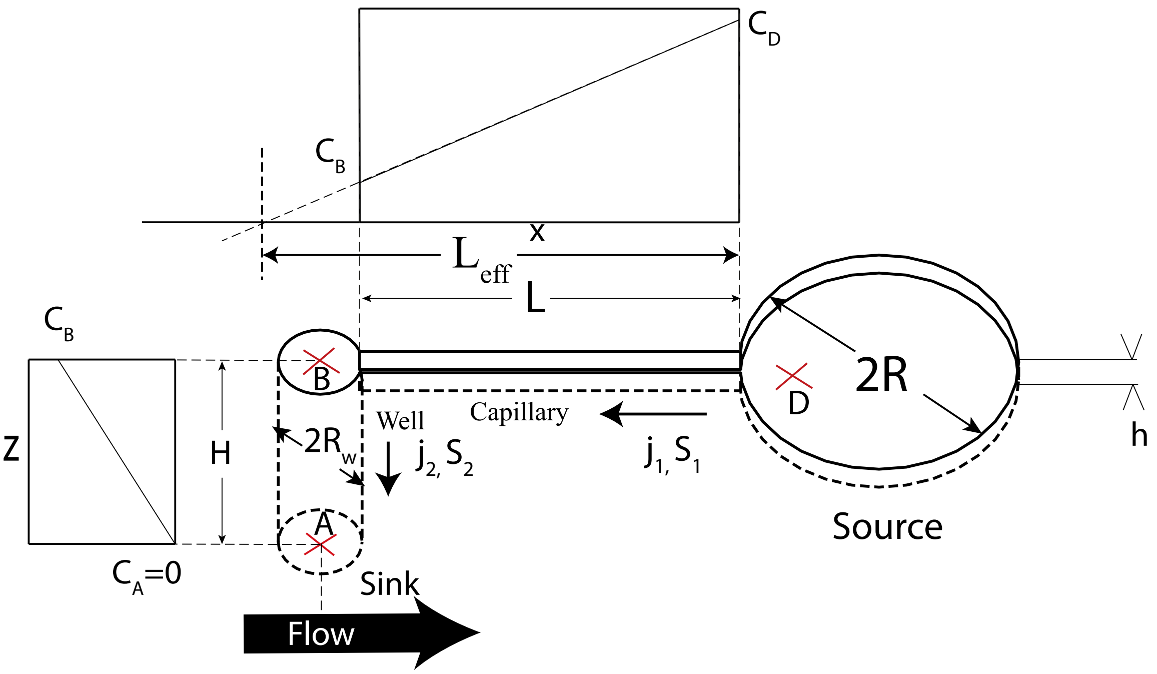


The compartment contains the DNA brush where the protein synthesis occurs. Homogenization inside the compartment is faster than the diffusion alongside the narrow capillary. Therefore, we consider the whole compartment as the source for protein and the concentration homogeneous. The end of the well which is in contact with the feeding microchannel is continuously replenished by new cell extract. The proteins diffuse to this end of the well are washed away by the flow, thus creating a sink for proteins.

The well which is wider than the narrow capillary adds up a contribution to the diffusive capillary connecting the well and the compartment[3]. We calculate the concentration of GFP produced in the compartment and diffusing out along the axis composed of the capillary and the well.

The exiting flux from the compartment is

$$J = j_1 S_1 = j_2 S_2 \tag{3.1}$$

$S_1$ and $S_2$ are the cross section of the capillary and the well respectively.
$j_1$ and $j_2$ are the diffusion flux (amount of substance per unit area per unit time) of the capillary and the well.
In permanent regime, the flux per surface unit is given by the Fick's Law

$$\vec{j} = -D_0 \overrightarrow{grad} C \tag{3.2}$$

3.2 integrated between the compartment D and the end of the capillary B gives:

$$j_1 = -\frac{D_0(C_B - C_D)}{L_1} \tag{3.3}$$

And between the flow channel A and the end of the well B

$$j_2 = -\frac{D_0(C_A - C_B)}{L_2} \tag{3.4}$$

By conservation of mass, we can write the equality of the fluxes:

$$-S_1.\frac{D(C_B - C_D)}{L} = -S_2.\frac{D(C_A - C_B)}{H} = -S_1.\frac{D(C_A - C_D)}{L_{eff}} \tag{3.5}$$

$L_{eff}$ is the length of the equivalent capillary that would connect A and D with the cross-section $S_1$.

$$\frac{L_{eff}}{L} = \frac{C_A - C_D}{C_B - C_D} = \frac{C_A - C_B + C_B - C_D}{C_B - C_D} = 1 + \frac{C_A - C_B}{C_B - C_D} = 1 + \frac{S_1 H}{S_2 L} \tag{3.6}$$

$$S_1 = W_c h, S_2 = \pi R_w^2$$

Therefore,

$$\frac{\mathrm{L_{eff}} - L}{L} = \frac{whH}{\pi R_w^2 L} \approx 12\% \tag{3.7}$$

By conservation of mass inside the compartment, the DNA brush creates at steady state $a_0$ proteins (in mol/s). Therefore we can write

$$\partial_t V C_D = a_0 + J \tag{3.8}$$

Replacing J,

$$\partial_t C_D = \frac{a_0}{V} - \frac{S_1 C_D D}{V L_{eff}} \tag{3.9}$$

At steady state:

$$0 = \frac{a_0}{V} - \frac{whDC_D}{\pi R^2 hL\left(1 + \frac{\Delta L}{L}\right)} \tag{3.10}$$

$$C_D = \frac{a_0 \tau_0}{V}\left(1 + \frac{\Delta L}{L}\right) \tag{3.11}$$

With $\tau_0 = \frac{\pi R^2 L}{wD}$

The observed extension (Figure S2) is about 18%. This is in the same order of the 12% calculated above. However, the model assumes as a first approximation that diffusion in the well is 1D.

### III. Flow between compartments, diffusion versus convection

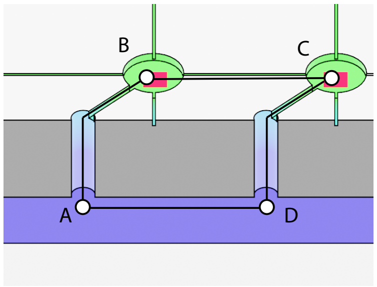

Let's estimate the flow between two compartments and compare it to diffusion
$P_X$ is the pressure at the position $X$. $R_{XY}$ is the hydrodynamic resistance between the positions $X$ and $Y$. $Q_{XY}$ is the flow in the branch $XY$.
The analogy of Ohm's law for hydrodynamics gives:

$$P_D - P_A = R_{AD} Q_{AD} \tag{4.1}$$

$$P_D - P_A = (R_{AB} + R_{BC} + R_{CD}) Q_{BC} \tag{4.2}$$

$$\frac{Q_{BC}}{Q_{AD}} = \frac{R_{AD}}{R_{AB} + R_{BC} + R_{CD}} = \frac{R_{AD}}{2R_{AB} + R_{BC}} \tag{4.3}$$

Let's write the expression for hydrodynamic resistances function of $\eta$, the dynamic viscosity of the cell-extract (square section):
$(L = 80\mu m, w = 10\mu m, h = 3\mu m, h_{backside} = 120\ \mu m, w_{backside} = 50\ \mu m)$

$$R_{AD} = \frac{12\eta L}{w_{backside} h_{backside}^3 \left(1 - \frac{0.63 h_{backside}}{w_{backside}}\right)} \approx 1.6.10^{-4}\eta \tag{4.4}$$

$$R_{AB} = \frac{12\eta L}{w h^3 \left(1 - \frac{0.63h}{w}\right)} \approx 5.5\eta \tag{4.5}$$

$$R_{BC} = \frac{12\eta L}{w h^3 \left(1 - \frac{0.63h}{w}\right)} \approx 5\eta \tag{4.6}$$

Therefore, the ratio of the fluxes is:

$$\frac{R_{AD}}{2R_{AB} + R_{BC}} \approx 1.10^{-5} \tag{4.7}$$

Estimation of the fluid velocity between B and C, from paragraph SI.2
$Q_{AB} = 0.01\mu L/min = 1.7.10^5 \mu m^3/s$ , $h = 3\mu m$, $w = 10\mu m$

$$V_{BC} = \frac{Q_{BC}}{hw} = 1.10^{-5} * \frac{Q_{AB}}{hw} \approx 0.05\ \mu m/s$$

Estimation of the time for a fluid particle to be transported from one compartment to another one ($L_c = 80\mu m$):

$$\tau_{conv} \approx 30\ min$$

Estimation of the time for a protein to diffuse from one compartment to another one:

$$\tau_{diff} = \frac{L_c^2}{D_0} \approx 3\ min$$

$$\tau_{conv} \gg \tau_{diff}$$

We conclude that diffusion is the main transport mechanism inside the array.

IV. Establishing the reaction-diffusion equation in a coupled array

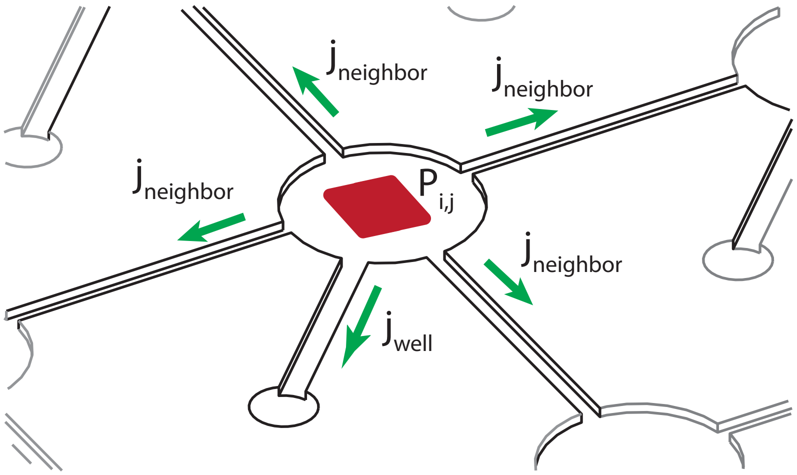


Exiting fluxes of the compartment (Fick's Law):

$$j_{well} = -D\frac{P}{l} \quad (5.1)$$

$$j_{neighbor} = -D\frac{P - P_{neighbor}}{L_c} \quad (5.2)$$

Conservation law gives

$$\partial_t V P_{i,j} = hw.j_{well} + hw_c \sum_{neighbors} j_{neighbor} + F(P_{i,j}) \quad (5.3)$$

$$\partial_t P_{i,j} = -\frac{hwD}{Vl}P_{i,j} - \frac{hw_C D a^2}{V l_C}\frac{4P_{i,j} - P_{i+1,j} - P_{i-1,j} - P_{i,j-1} - P_{i,j+1}}{a^2} + f(P_{i,j}) \quad (5.4)$$

F represents the production rate of the protein P given the concentration of P inside the compartment, f is the same function per unit volume (SI.VI).

We recognize here

$$\tau_0 = \frac{Vl}{hwD} = \frac{\pi R^2 l}{w D_0} \quad (5.5)$$

And

$$\tau_c = \frac{V l_C}{h w_C D} = \frac{\pi R^2 l_C}{w_C D_0} \quad (5.6)$$

Thus

$$\partial_t P_{i,j} = -\frac{1}{\tau_0}P_{i,j} - \frac{a^2}{\tau_C}\frac{4P_{i,j} - P_{i+1,j} - P_{i-1,j} - P_{i,j-1} - P_{i,j+1}}{a^2} + f(P_{i,j}) \quad (5.7)$$

Finally

$$\partial_t P_{i,j} = -\frac{P_{i,j}}{\tau_0} + \frac{a^2\beta}{\tau_0}\Delta P_{i,j} + f(P_{i,j}) \quad (5.8)$$

With $\beta = \tau_0/\tau_c$ and $\Delta P_{i,j} = \frac{P_{i+1,j}+P_{i-1,j}+P_{i,j-1}+P_{i,j+1}-4P_{i,j}}{a^2}$

In the continuous limit:

$$\partial_t P = -\frac{1}{\tau_0}P + \frac{a^2\beta}{\tau_0}\Delta P + f(P_{i,j}) \quad (5.9)$$

$$D_{eff} = \frac{a^2\beta}{\tau_0}$$

The dimensionless equation is:

$$\boxed{\partial_t P = -P + \Delta P + f_0(P_{i,j}) \quad (5.10)}$$

With $t' = t/\tau_0$ , $x_i' = x_i/a\sqrt{\beta}$ and $f_0 = \tau_0 f$

Single source analytical solution

Let's solve the problem of diffusion from a single steady source placed at $\vec{r} = \vec{0}$ producing $a_0$ in mol/s

The protein concentratTapez une équation ici.ion does not depend on $\theta$

The previous equation becomes in steady state for $r \neq 0$:

$$\frac{1}{r}\frac{dP}{dr} + \frac{d^2P}{dr^2} - P = 0 \quad (5.11)$$

$$r\frac{dP}{dr} + r^2\frac{d^2P}{dr^2} - r^2P = 0 \quad (5.12)$$

This equation is the modified Bessel equation, and its solutions are a linear combination of $I_0$ and $K_0$. $I_0$ does not correspond to a physical solution because it is diverging for $r \to \infty$.

At $r = R_0$the outcoming flux is:

$$0 = a_0 - \frac{P(R_0)}{\tau_0} - \frac{\beta}{\tau_0}\oint\left(-\frac{dP}{dr}\right)rd\theta \quad (5.13)$$

$$a_0\tau_0 = P - \beta 2\pi R_0\frac{dP}{dr} \quad (5.14)$$

Continuity in $R_0$

$P(r) = \lambda K_0(r)$ and $\frac{dP}{dr} = \lambda K'_0(r)$

$$a\tau_0 = \lambda K_0(R_0) - 2\pi R_0\beta\lambda K'_0(R_0) \quad (5.15)$$

$$\lambda = \frac{a_0\tau_0}{K_0(R_0) - 2\pi R_0 K'_0(R_0)} \quad (5.16)$$

$\lambda$ is a parameter that depends on the cell-extract and the DNA brush. It can be adjusted to fit the experimental data.

V. Model for the genetic oscillator

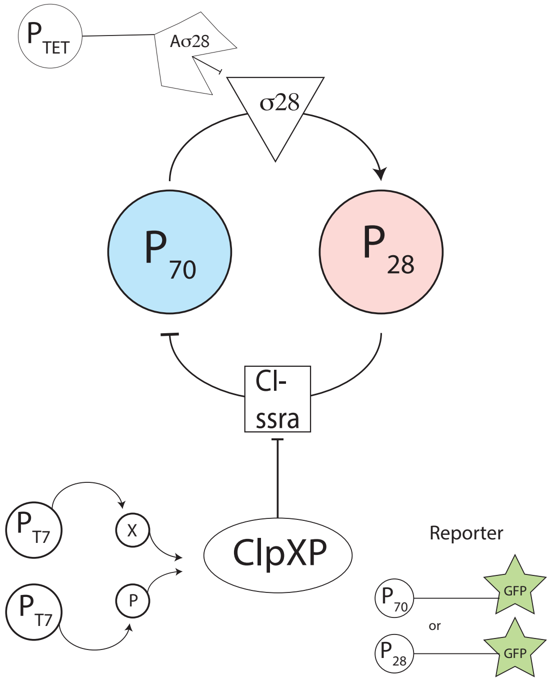


Our set of 5 genes generates oscillations in the compartments. The activator is the protein σ28 and the inhibitor is the protein CI-ssra They are made from the genes (P70-*σ28* and P28-*ci-ssra*). These two proteins are degraded respectively by the protein Anti-σ28 and ClpXP (Respectively from the genes pTET-*Aσ28*, PT7-*clpX* and PT7-*ClpP*). The concentration of σ28 and CI-ssra are reported alternatively by the gene P70-*gfp* or by P28-*gfp*.
For the simulations, we use a set of 4 partial differential equations[4] for the concentrations of the activator (A, σ28), repressor (R, CI) and their corresponding mRNAs ($m_A$ and $m_R$).
Oscillations are spontaneously generated by these equations in single compartments ($\beta = 0$). Their period scales linearly with $\tau_0$.

$$\frac{dm_A}{dt} = k_{TX} D_A \frac{1}{1 + \frac{R_i^4}{K_{CI}}} - \frac{m_A}{\tau_m} \quad (6.1)$$

$$\frac{dA}{dt} = k_{TL} m_A - \frac{A}{\tau_0} + \frac{a^2 \beta}{\tau_0} \Delta A_{i,j} \quad (6.2)$$

$$\frac{dm_R}{dt} = k_{TX} D_R \frac{A}{A + K_{28}} - \frac{m_R}{\tau_m} \quad (6.3)$$

$$\frac{dR}{dt} = k_{TL} m_R - \frac{R}{\tau_0} - C\Theta(R - R_0^*) + \frac{a^2 \beta}{\tau_0} \Delta R_{i,j} \quad (6.4)$$

$k_{TX}$ and $k_{TL}$ are transcription and translation rates of the mRNA and proteins respectively.
$D_A$ and $D_R$ are the DNA concentrations of the activator and repressor genes in the DNA brush
$K_{CI}$ and $K_{28}$ are the Michaelis-Menten constants for binding between σ28 and P28 and CI to P70

$\tau_m$ is the mRNA lifetime in the comapartment
C is the degradation rate of ClpXP
$R_0^*$ is the concentration threshold tuning on the activity of ClpXP
Simulations with the following parameters are plotted in Figure S4e :

| **Parameter** | **Value** |
|---|---|
| $k_{TX}$ | $0.15\ min^{-1}$ |
| $k_{TL}$ | $1\ min^{-1}$ |
| $K_{CI}$ | $1\ nM^4$ |
| $K_{28}$ | $0.7\ nM$ |
| $\tau_m$ | $10\ min$ |
| C | $50\ nM/min$ |
| $R_0^*$ | $1\ nM$ |

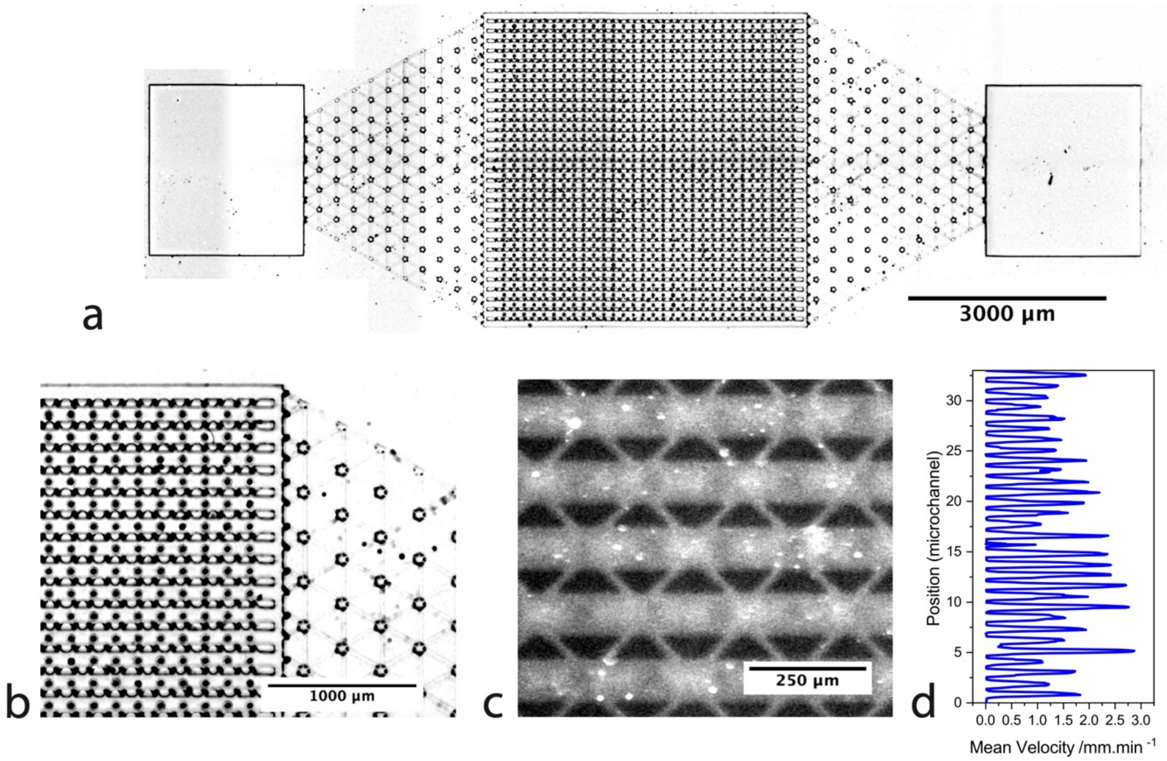


**Fig. S1: Backside of the chip and parallel channels.**

A. Micrograph of the backside of the chip (Scale bar 3 mm). Inlet are pierced through the sealing PDMS slab. Cell-free extract flow is then divided into 32 parallel flow-channels. Every channel is connected to 32 wells, connecting the compartments and the flow channels. B Close up on the parallel microchannels and the flux divider (scale bar 1000 µm). C. Fluorescent particles in the backside flow-channel. Measuring their speed give provides the mean speed in the flow-channels. (scale bar 250 µm). D Mean velocity of the cell-free extract measured for every channel by PIV (PIVlab)

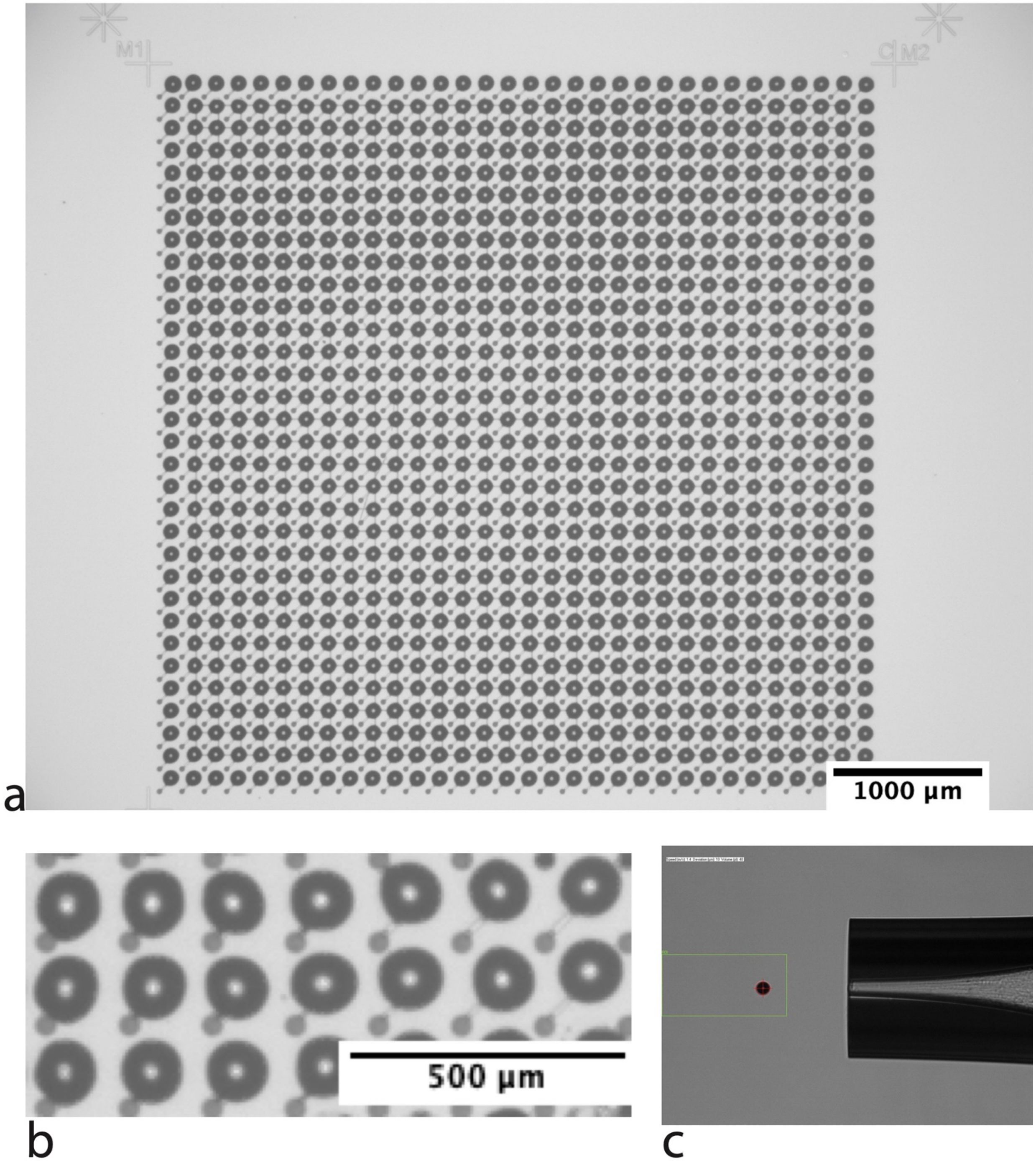


**Fig. S2: Spotting DNA solutions**

A. Micrograph of the chip covered with 30 pL droplets. Each droplet was independently dispensed. Scale bar 1000 µm B. Close up on dispensed droplets. Compartments have varying compartment lifetimes. Scale bar 5000 µm C. Image of the tip expelling a droplet of 40 pL.

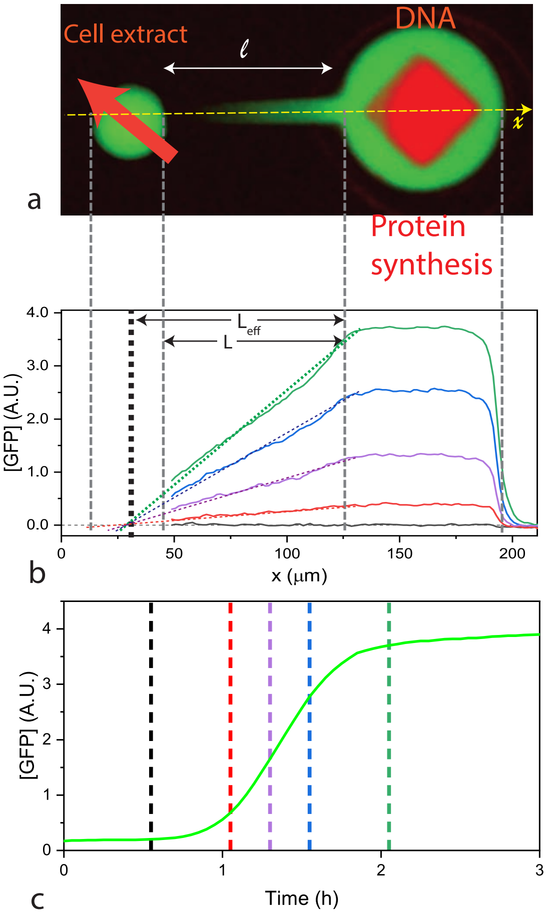


**Fig. S3: Expression concentration gradients in a single artificial cell in a 3D geometry.**
A. Epifluorescence image of a compartment during protein synthesis: end-labeled DNA brush (red) and GFP (green). B. GFP expression spatial profiles at different time points as denoted. The linear profiles at different time points extrapolate to a point inside the well where [GFP]=0. $L = 80\ \mu m$, $L_{eff} = 95\ \mu m$. The extension of the linear GFP profile inside the capillary meets the zero baseline at $x = 30 - 35\ \mu m$. This point corresponds to an estimated $\frac{\Delta L}{L} = 18\%$ extension of the capillary length. C. Dynamics of GFP expression from onset to steady-state, the colored dashlines indicate the time for which the profile were extracted in B.

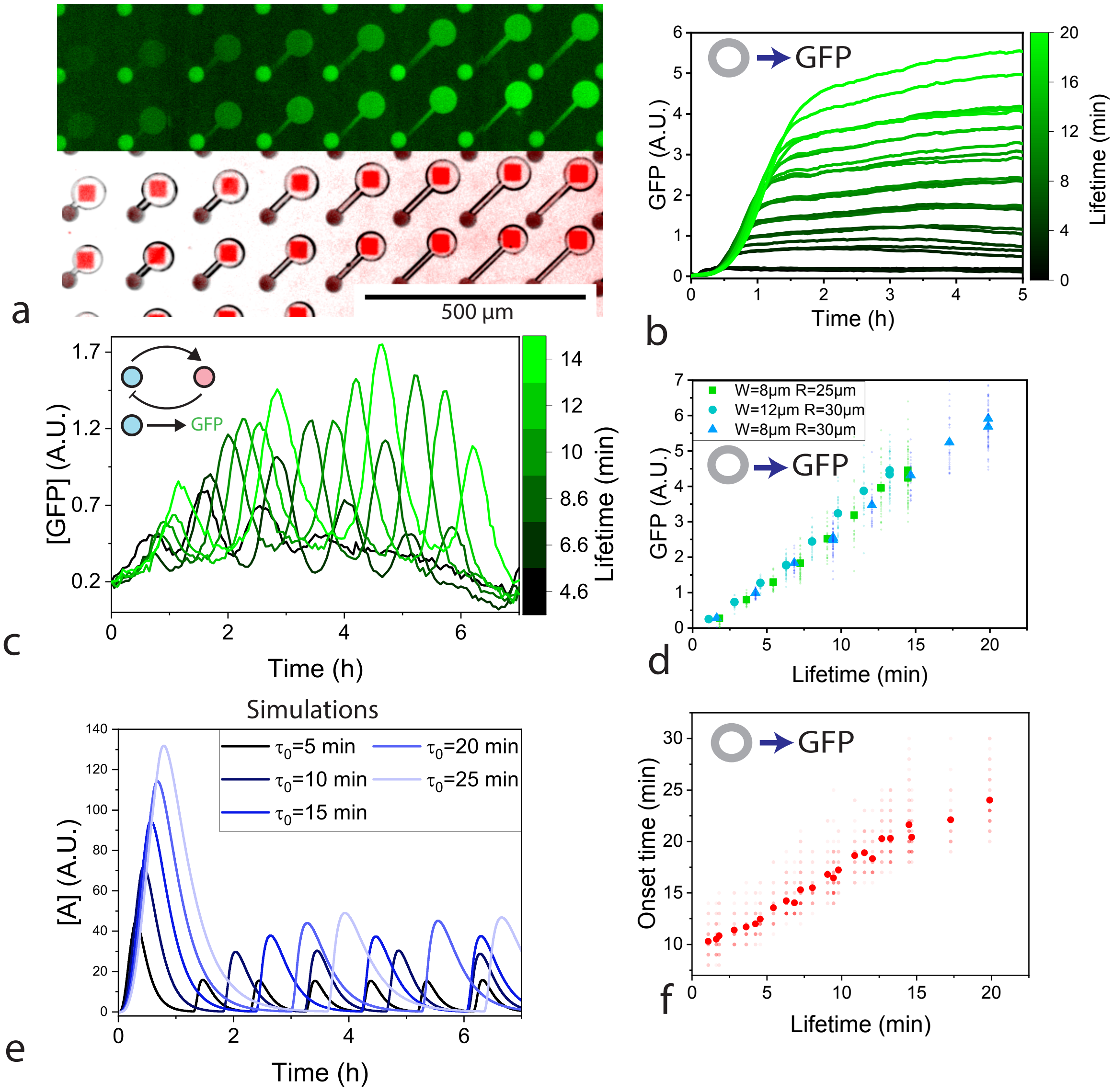


**Fig. S4 Effect of the lifetime on GFP expression, oscillations, and onset times.**

A. Left: Fluorescent signal of GFP expression (Top, green) from DNA brushes (Bottom, end-labelled, red squares) at varying compartment radius, and capillary width and length, setting the lifetime from 2 to 20 min. Scale bar 500 µm. B. Dynamics of GFP expression as a function of lifetime under P70-GFP gene construct. C,E. GFP expression (C) under a genetic oscillator at varying compartment lifetime. Simulated oscillations for different lifetimes (E) D,F. Steady-state concentration (D) and onset time (F). The fit crosses the y-axes at y=8min, possibly related to the maturation time of GFP in cell-extract.

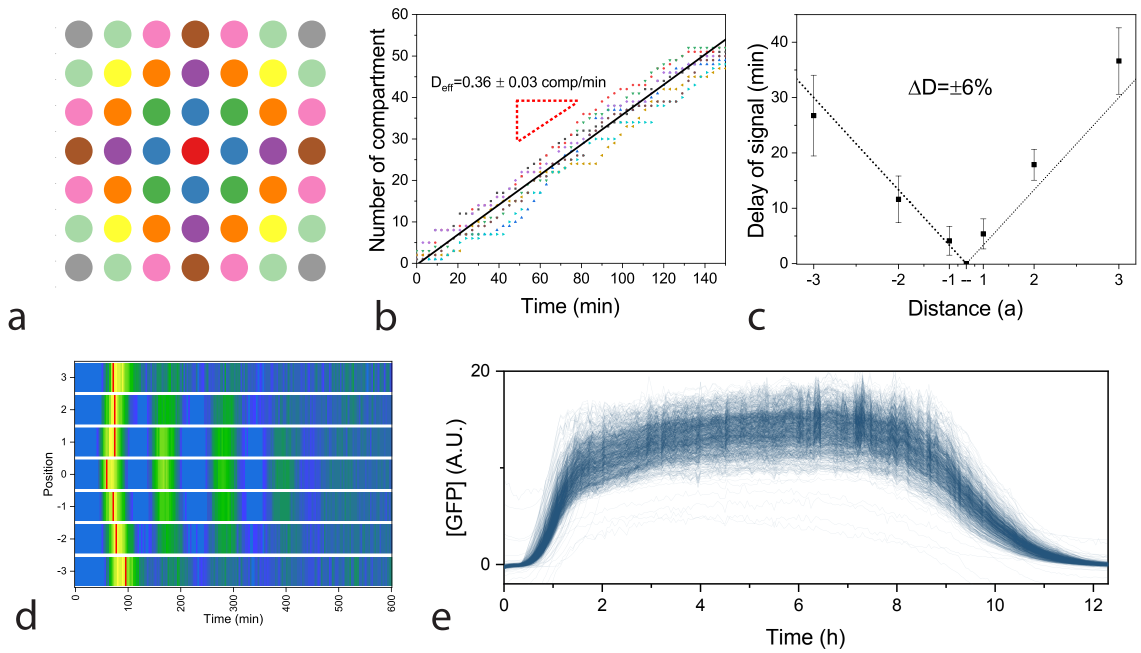


**Fig. S5: Diffusion of expressed protein from a localized source in a coupled array**

A. Color code for main text Figure 2. B. Cumulated area of diffusion from a single source in time in an array of coupling strength $\beta = 1.1$. The linear fit implies an effective diffusion constant of $D_{eff} = 27 \pm 2\ \mu m^2/s$ . The surface unit for compartments is $\pi R^2 + 2w_c L_c = 4500\mu m^2$ C. The delay in signal of diffusing proteins as a function of distance from source in the x-direction. The asymmetry of 6% comes from a minor drift due to the flow at the backside from left and right. D. Space-time plot of a single oscillator diffusing into neighboring compartments on the same array. Delays between two compartments is measured at $\Delta t = 9\ min$ E. Expression of GFP under the P70-GFP construct in an array of 30x30 coupled compartments.

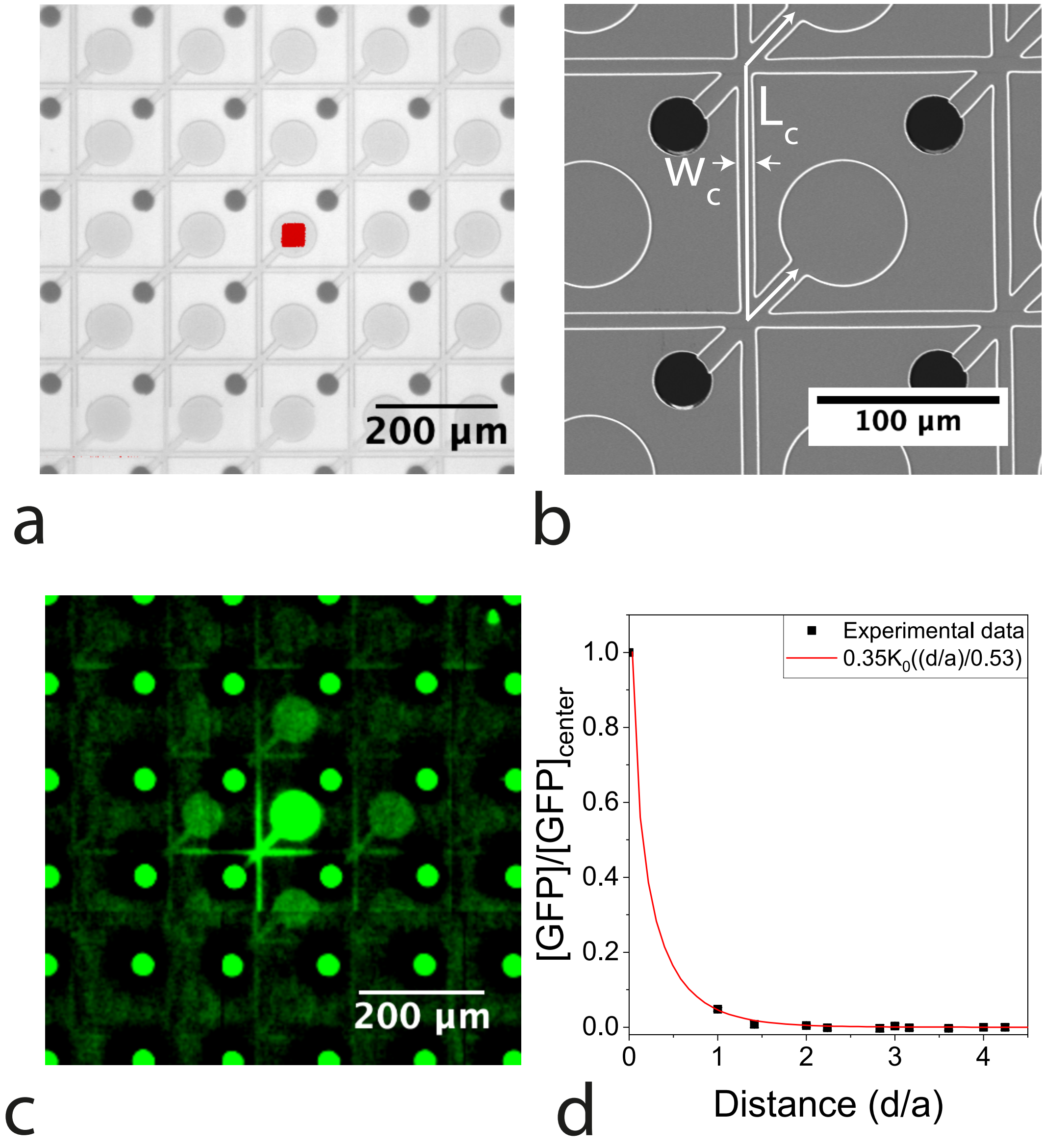


**Fig. S6: Diffusion of expressed protein from a localized source in a coupled array – Low coupling.**

A Micrograph of the chip ($\beta = 0.3$). Scale bar 200µm. DNA signal is overlayed and reveal the single source of GFP placed at the center of the image. B Electron micrograph of the compartment loosely coupled to its neighbors. $w_c = 8\ \mu m$ and $L_{c,eq} = 220\ \mu m$. Scale bar 100µm. C Fluorescent signal of GFP. A single source is present at the center and GFP diffuses out to the neighboring compartments through the coupling capillaries. D Profile of the GFP expression with the distance to the source. The signal was normalized to the source. The GFP concentration is fit by the function $f(d) = \lambda K_0(\frac{d}{a\sqrt{\beta}})$. Effective diffusion constant is estimated at $D_{eff} = 10\ \mu m^2/s$.

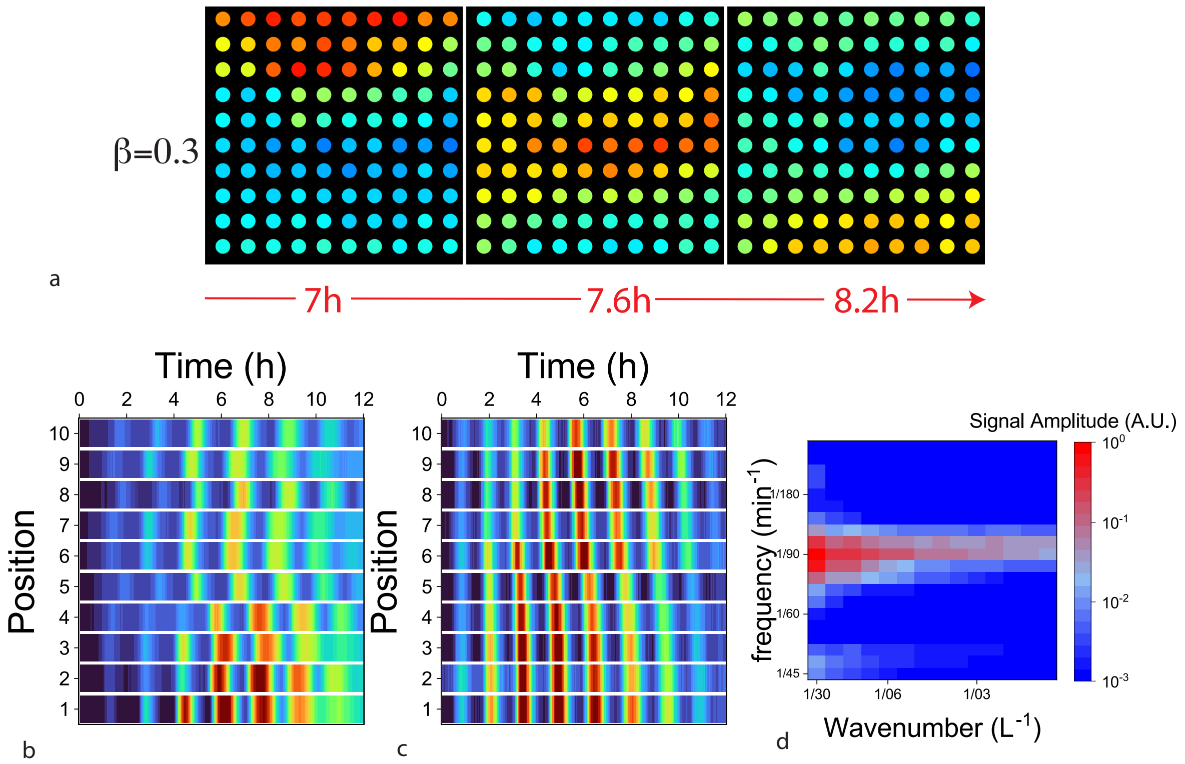


**Fig. S7: Wave Propagation for different coupling strength.**

A Close-up on a 10x10 window of coupled genetic oscillators expressed in an array with coupling strength of $\beta = 0.3$. A phase wave travelling from the top to the bottom of the window, shown at three time points, as denoted. B Space-time plot for 10 neighboring compartments that are not coupled ($\beta = 0$). C Space-time plot for 10 consecutive oscillating compartments that are weakly coupled ($\beta = 0.3$) D. Power spectrum of the weakly coupled system ($\beta = 0.3$).

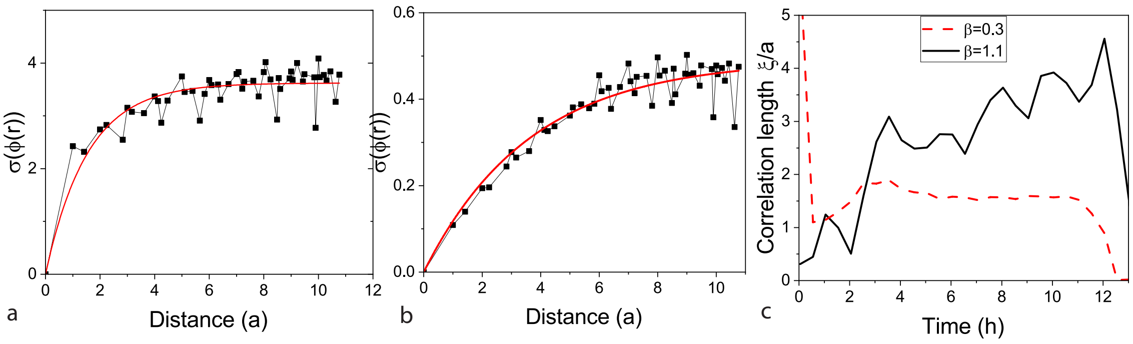


**Fig. S8: Correlation length**

A-B Standard deviation of the phase at $t = 10\,h$ for coupling strength $\beta = 0.3$ (A) and $\beta = 1.1$ $(B)$ plotted against the size of the considered window R in unit $a$, with a fit to an exponential function (solid red line), yielding a correlation length of $\xi = 1.6a$, and $\xi = 3.9a$, respectively. C The correlation length plotted with time as x-axis for $\beta = 1.1$ and $\beta = 0.3$.

**Movie S1.**

Experimental movie corresponding to single sources in a 30x30 compartment array.

**Movie S2.**

Experimental movie corresponding to 30x30 oscillating compartments


1. Buxboim, A. *et al.* A Single-Step Photolithographic Interface for Cell-Free Gene Expression and Active Biochips. *Small* **3**, 500–510 (2007).
2. Garamella, J., Marshall, R., Rustad, M. & Noireaux, V. The All E. coli TX-TL Toolbox 2.0: A Platform for Cell-Free Synthetic Biology. *ACS Synth Biol* **5**, 344–355 (2016).
3. Karzbrun, E., Tayar, A. M., Noireaux, V. & Bar-Ziv, R. H. Programmable on-chip DNA compartments as artificial cells. *Science (1979)* **345**, 829–832 (2014).
4. Tayar, A. M., Karzbrun, E., Noireaux, V. & Bar-Ziv, R. H. Synchrony and pattern formation of coupled genetic oscillators on a chip of artificial cells. *Proceedings of the National Academy of Sciences* **114**, 11609–11614 (2017).
